# Development of metastases in mice induced by plasma sample collected during radiotherapy in patient with triple-negative breast cancer: Role of Rab4A

**DOI:** 10.1101/2025.07.23.666270

**Authors:** Benoit Paquette, Hélène Therriault, Isabelle Gauthier, Sawyna Provencher, Sahar Naasri, Ayman Oweida, Sameh Geha, Pascale Labrecque, Jean-Luc Parent

**Affiliations:** Centre for Research in Radiotherapy, Sherbrooke University Cancer Research Institute, Department of Medical Imaging and Radiation Sciences, University of Sherbrooke, Sherbrooke, Quebec, Canada; Service of Radiation Oncology, Sherbrooke University Hospital Center, Sherbrooke, Quebec, Canada; Department of Pathology, Sherbrooke University Hospital Center, Sherbrooke University Cancer Research Institute, Universite de Sherbrooke, Sherbrooke, Quebec, Canada; Department of Medicine, Faculty of Medicine and Health Sciences, University of Sherbrooke, Sherbrooke, Quebec, Canada; Sherbrooke Institute of Pharmacology, Faculty of Medicine and Health Sciences, University of Sherbrooke, Sherbrooke, Quebec, Canada

## Abstract

The relapse rate in early-stage triple-negative breast cancer (TNBC) is significantly higher than in other breast cancer subtypes. This study assessed the relevance to target *RAB4A* to prevent the development of metastases that occur after treatment. The ability of cancer cells to invade peritumoral tissue is associated with the expression of membrane-type matrix metalloproteinase-1 on their surface, which is regulated by RAB4A. When RAB4A was downregulated using shRNA in the TNBC cells D2A1 and MDA-MB-231, a significant reduction in the proteolytic activity of MT1-MMP and the invasion capacity of these TNBC cells were measured. Plasma samples from an early-stage TNBC patient, who developed metastases six months after treatment, were collected before radiotherapy and after the fourth radiation dose. Compared to the plasma collected before radiotherapy, the plasma collected during the treatment significantly enhanced the invasiveness of the TNBC cells, as assessed with Boyden chambers. The development of lung metastases was also stimulated when the D2A1 cells were preincubated with this plasma before their i.v. injection in female Balb/c mice. Importantly, these adverse effects of plasma collected during radiotherapy were significantly blocked by downregulating RAB4A. These results highlight the relevance of developing RAB4A inhibitors to prevent the development of metastases occurring after treatment in TNBC patients.

## Introduction

The triple negative breast cancer (TNBC) subtype is characterized by low or absent expression of estrogen, progesterone and human epidermal growth factor-2 (HER2) receptors [1]. Although radiotherapy improves survival in patients with breast cancer [2], the risk of recurrence remains higher in those with TNBC [3]. The recurrence pattern is also different. It occurs in ∼30% of them, with risk peaking at ∼3 years after treatment [3, 4].

Cancer cells can leave the primary tumor and infiltrate the breast [5]. In order to treat them, the whole breast is irradiated as well as certain locoregional lymph nodes [6]. A total dose of 42.5 to 50 Gy is delivered in 16 to 25 fractions [7]. Therefore, the first fractions of radiation are not lethal to a large proportion of cancer cells [8]. Moreover, the total dose that can be delivered is limited by the tolerance of the normal tissue included in the irradiated volume [9]. Therefore, a lethal dose of radiation may not be delivered to all cancer cells, which could contribute to local recurrence and development of metastases [10].

Metastases detected after treatment may originate from micrometastases present at diagnosis but undetectable [11], or may be induced by a sublethal dose of radiation [11–13]. Indeed, clinical and preclinical studies suggest that radiotherapy induces changes in the tumor environment that may increase the risk of developing metastases and thus diminish the long-term effectiveness of the treatment [8, 14–18].

Pre-irradiation of the mouse mammary gland followed by implantation of non-irradiated TNBC cells, or irradiation of a TNBC tumor implanted in a mammary gland has been shown to increase cancer cell invasion in the mammary gland and the number of circulating tumor cells, which has resulted in a greater number of lung metastases [19, 20]. These adverse effects of radiotherapy were blocked by reducing expression of the membrane-type matrix metalloproteinase-1 (MT1-MMP) on the TNBC cells [21]. These results indicate that the higher number of circulating tumor cells and lung metastases was not significantly caused by an increase in vascular permeability but rather by a stimulation of the invasive capacity of cancer cells [22].

The expression of MT1-MMP on the surface of cancer cells plays a central role in the invasion process. Its upregulation is associated with malignancy in several types of cancer, including for those of breast [23]. This transmembrane protease can activate the matrix metalloproteinases-2 and -9 (MMP-2 and MMP-9) [24, 25], and in coordination with them cleave extracellular matrix proteins and thus open a passage for cancer cells [26, 27]. In TNBC patients, elevated levels of MT1-MMP have been correlated with cancer cell invasion of blood vessels [28], and shorter overall survival [29].

Endocytic and exocytic cycles of MT1-MMP depend on Rab4A [23, 30]. Rab4A is a member of the Rab GTPases family that coordinates vesicle traffic [31, 32]. The importance of Rab4A in invasion and metastasis formation is supported by the progressive increase of its expression from normal tissue to primary tumor and then to metastases [33]. Rab4A expression is amplified in various tumors and frequently in invasive breast carcinoma [33, 34]. RAB4A could therefore be a good target to prevent radiation-stimulation of metastasis development.

In the present study, the role of Rab4A in radiation-stimulation of TNBC cell invasion *in vitro* and the development of metastases in animal model was determined. In order to get closer to the clinic, plasma samples from a TNBC patient were collected before radiotherapy and after the 4^th^ radiation fraction. This patient was selected because fluorodeoxyglucose-positron emission tomography (FDG-PET) imaging performed prior treatment did not detect cancer cells in the lymph nodes, nor the presence of metastases. Unfortunately, metastases in lung, bone and brain appeared six months after radiotherapy. This patient represents an example of the rapid recurrence observed in approximately 30% of women with TNBC. The TNBC cells D2A1 (murine) and MDA-MB-231 (human) were treated with these plasmas and we determined if the downregulation of RAB4A prevents the stimulation of cancer cell invasion and metastasis development that were induced by the plasma collected during radiotherapy.

## Materials and Methods

### Cell culture

The mouse breast carcinoma D2A1 cells derived from a spontaneous mammary tumour in a Balb/c mouse and were provided by Dr Ann F. Chambers (University of Western Ontario, London, ON, Canada). The human MDA-MB-231 cells were purchased from American Type Culture Collection. The cells were maintained in a 5% CO_2_ humidified incubator at 37°C in Dulbecco’s modified Eagle’s medium (DMEM) supplemented with 10% fetal bovine serum, 2 mM glutamine, 1 mM sodium pyruvate, 100 units per mL penicillin and 100 mM streptomycin. All experiments were performed with mycoplasma-free cells. Cells were tested for mycoplasma by real time qPCR coupled to capillary electrophoresis at the RNomics Platform of the Université de Sherbrooke (https://rnomics.med.usherbrooke.ca/services/mycoplasma-detection).

### Downregulation of RAB4A and MT1-MMP expression

A short hairpin RNA (shRNA) mediated gene knockdown was performed as previously reported [21]. Lentiviral particles were produced in the human cell line 293T (5x10^6^) which was co-transfected using 48 µg lipofectamine with 6 mg of the plp1, plp2 and plp/VSV-G plasmids (Invitrogen) and 6 mg of pLKO.1-puro vector containing a shRNA sequence targeting either the murine RAB4A transcript TRCN0000088973, TRCN0000088974, TRCN0000088975, TRCN0000088976 and TRCN0000088977, or the human RAB4A transcript TRCN0000011217, TRCN0000231999 and TRCN0000232000 (Sigma-Aldrich). As control, the PLKO.1 non-target control DNA (code SHC016) was used. The cultured cell supernatant containing the lentivirus was collected 48 h later, filtered with a 0.45 µm membrane and kept at -80°C for further use.

The downregulation of MT1-MMP expression by shRNA in D2A1 cells was previously reported [21]. A 70% reduction of MT1-MMP mRNA was confirmed by quantitative polymerase chain reaction (qPCR) and at the protein level by Western blot. The ability of MT1-MMP to cleave inactive proMMP-2 to active MMP-2 in D2A1 MT1-MMP knockdown cells was reduced by more than 90% compared to wild-type D2A1 cells. Nomenclature of the derived D2A1 cell lines is as follows: D2A1-WT (wild-type), D2A1 RAB4A-scramble, D2A1 KD RAB4A (RAB4A downregulated) and D2A1 KD MT1-MMP (downregulation of the Mmp14 transcript level) cells. For those derived from the MDA-BM-231 cells: MDA-MB-231-WT, MDA-MB-231 RAB4A-scramble, MDA-MB-231 KD RAB4A.

### Western blot analysis

Down regulation of RAB4A was confirmed by Western blot. Briefly, cells were plated in 100 mm plates and harvested 24h later in in 600µl lysis buffer (150 mM NaCl, 50 mM Tris-HCl, pH 8.0, 0.5% deoxycholate, 0.1% SDS, 10 mM Na4P2O7, 1% IGEPAL, and 5 mM EDTA) supplemented with protease inhibitors (10 µM chymostatin, 10 µM leupeptin, 9 µM antipain and 9 µM pepstatin). Sample buffer was added to each reaction before boiling tubes for 5 minutes. All reactions were analyzed by Western blot (30 µg proteins/lane) using the anti-Rab4A (Santa Cruz Biotechnology # sc-517263) and anti-actin (C4) (Santa Cruz Biotechnology) antibodies. Experiments were done 3 times and densitometry analyses were carried out using NIH Image J software.

### Analysis of the proteolytic activity of MT1-MMP

A fluorogenic peptide was used to measure the proteolytic activity of MT1-MMP on cell surface [35]. The D2A1 or MDA-MB-231 cells (3 x 10^4^ in 96-well plate) were incubated at 37°C with 15 µM of the fluorescence resonance energy transfer (FRET) peptide MMP-14 Substrate I (Sigma 444258-Calbiochem) in the reaction buffer (100 mM TRIS, 5 mM CaCl_2_, 0.01% BRIJ-35, 1 µM ZnCl_2_). The enzymatic activity was recorded according to the variation of relative fluorescent units per second (RFU/sec). The kinetics of peptide cleavage were followed for 30 min using the 96-well plate reader Synergy HT (Bio-Tek Instrument) set at λex = 340 nm and at λem = 400 nm.

### Collection of plasmas from TNBC patient

The research protocol was approved by the Research Ethics Committee, CIUSSS de l’Estrie CHUS, Quebec, Canada (protocol # MP-31-2015-930, 14-205). A 48-year-old woman with pathology-confirmed TNBC status was recruited on February 12, 2021. The information and consent form was presented and explained to her by the treating radiation oncologist, and the participant signed it in her presence. The clinical staging was cT2 N0 M0, grade 3 tumor with a diameter of 3 cm, no distant metastasis at diagnosis as determined by FDG-PET/CT scan (Figure 1A), BRCA1 and BRCA2 normal, Ki67 ˃ 95%. Neo-adjuvant chemotherapy with 4 cycles of adriamycin-cyclophosphamide followed by 6 cycles of paclitaxel and 2 cycles of carboplatin and paclitaxel was administered. After partial mastectomy and a biopsy of the sentinel lymph nodes, the final staging was ypT2 N0 with negative margin. Adjuvant radiotherapy consisted of 40 Gy in 15 fractions followed by a boost on the surgical cavity of 10 Gy in 4 fractions. A grade 2 radiodermatitis was observed. Metastases were detected by CT angiogram six months after radiotherapy in the lung as shown in Figure 1B.

**FIGURE 1.**
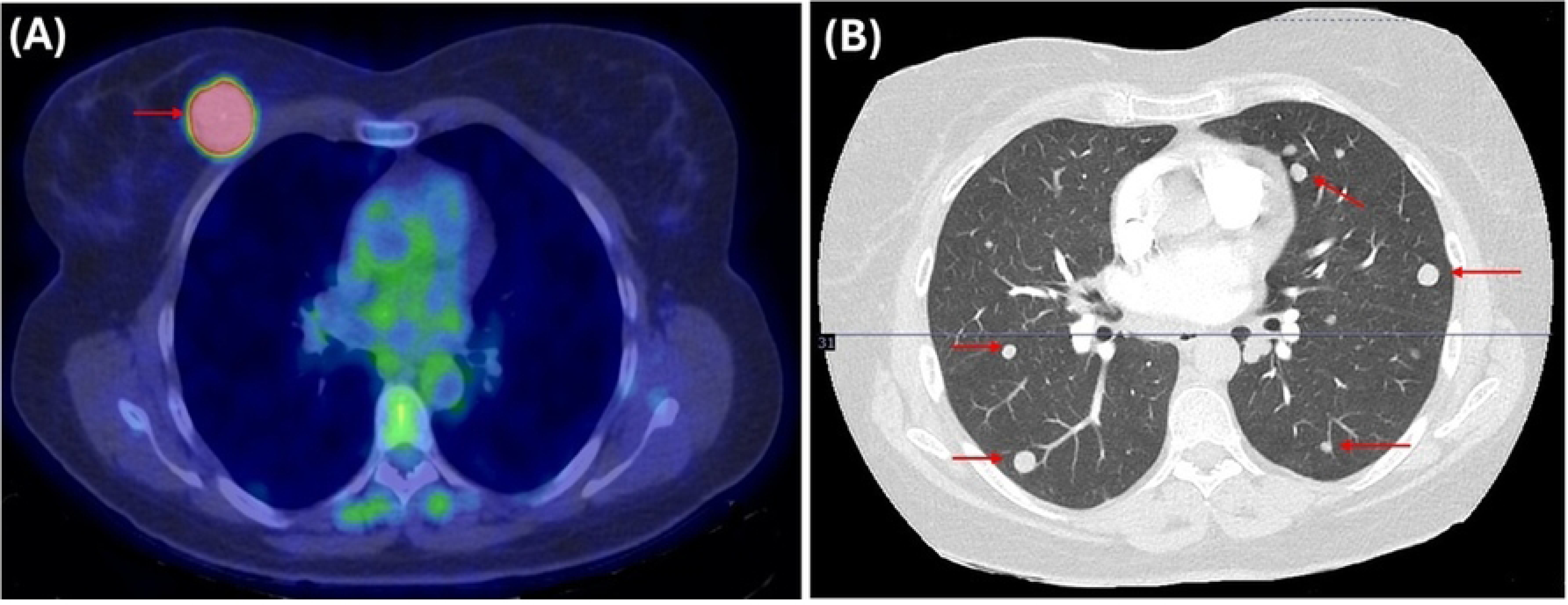
(A) Primary tumor located in a breast of a TNBC patient detected by positron emission tomography / computed tomography (PET/CT) imaging scan performed with the radiotracer [F18]-fludeoxyglucose (FDG) before any treatment. (B) Lung metastases detected six months after radiotherapy by CT angiogram imaging. Red arrows indicate the location of primary tumor and metastases.

A first blood sample was taken in an EDTA tube just before the first fraction of radiation (plasma before radiotherapy) and the second after the 4th fraction of radiation (plasma during radiotherapy). Plasmas were isolated by centrifugation at 1 200 g for 30 min within 40 min of collection. Aliquots of plasma were stored at −80°C.

### Cancer cell invasion assay

A volume of 100 µL of Matrigel (growth factor reduced, Corning) diluted 1/40 in cold DMEM 0.1% BSA was added on the porous membrane of cell culture insert (cellQART, Sterlitech) that was deposited in 24-well plate, and incubated for 2 h at 37°C to allow the polymerization of Matrigel. The liquid layer on the polymerized Matrigel was then removed. The TNBC cells D2A1 or MDA-MB-231 and their derivatives were incubated in 0.1% BSA DMEM medium for 24 h and then added (2 x 10^4^) on the Matrigel layer, while the lower compartment of the chamber contained either 1% plasma before radiotherapy, 1% plasma during radiotherapy, 1% FBS or 0.1% BSA in DMEM media. After 24 h at 37°C, cells on the top of the porous membrane were removed with cotton swabs while those on the bottom surface were stained with 0.4% crystal violet and then counted under a microscope. The ability of plasma collected during radiation to increase cancer cell invasion was reported by the number of cells that have crossed the layer of Matrigel in presence of 1% plasma collected during radiotherapy / 1% plasma collected before radiotherapy.

### Quantification of lung metastases

The experimental protocol in mice (Ref. # 2018-1996, 013-18) was approved by the ethical committee of Université de Sherbrooke and was conformed to the regulations of the Canadian Council on Animal Care. For the welfare of the mice and to minimize their suffering and distress, monitoring and care were carried out in accordance with the Canadian Council on Animal Care recommendations. The D2A1-WT, D2A1 RAB4A-scramble, D2A1 KD RAB4A cells were transfected with the fluorescent proteins FUCCI as previously reported to detect the lung metastases in mice by optical imaging [36]. These D2A1 cells were incubated during 24 h at 37°C in DMEM medium supplemented with either 1% plasma collected before radiotherapy, 1% plasma collected during radiotherapy, or in DMEM 0.1% BSA. The cells were washed twice by centrifugation in PBS and then injected (10^5^) in the tail vein of five female Balb/c mice (Charles River) per group for a total of 15 mice. The mice were monitored and weighed every 2 to 3 days to ensure that the development of lung metastases was not causing discomfort. Twenty-eight days after cell injection, the mice were euthanized with slow exposure of CO_2_, the lungs were collected, and the number of metastases was quantified with the epifluorescence microscope EVOS FL Auto Imaging System (Life Technologies) equipped with green (470/525 nm) or red (531/593 nm) light cubes. No mouse died before meeting the end point. Lung slices were stained by H&E and the presence of metastases were confirmed by a pathologist. Research staff were trained by an animal health technician at the animal facility for intravenous injection.

### Statistical analysis

Results are expressed as the mean ± standard deviation of 2 to 4 experiments performed in triplicate, or 5 mice regarding the metastasis assay. Statistical analyses were performed with PRISM v4.0 (GraphPad Software) using the Student’s t-test and two ways analysis of variance (ANOVA). A value of *p* < 0.05 was considered to be statistically significant. \**p* < 0.05, \*\**p* < 0.005, \*\*\**p* < 0.0005 and \*\*\*\**p* < 0.0005.

## Results

### Downregulation of RAB4A expression reduces proteolytic activity of MT1-MMP

The downregulation by shRNA of RAB4A expression in the D2A1 (murine) and MDA-MB-231 (human) cells was confirmed by Western blot analysis (Figure 2A and B). The level of Rab4A protein was more than 95% lower in the clone D2A1 KD RAB4A (shRNA sequence TRCN0000088975) and the clone MDA-MB-231 KD RAB4A (shRNA sequence TRCN0000232000), compared to their respective wild-type cells and shRNA scramble derivatives. These two clones were used for the subsequent assays.

**FIGURE 2.**
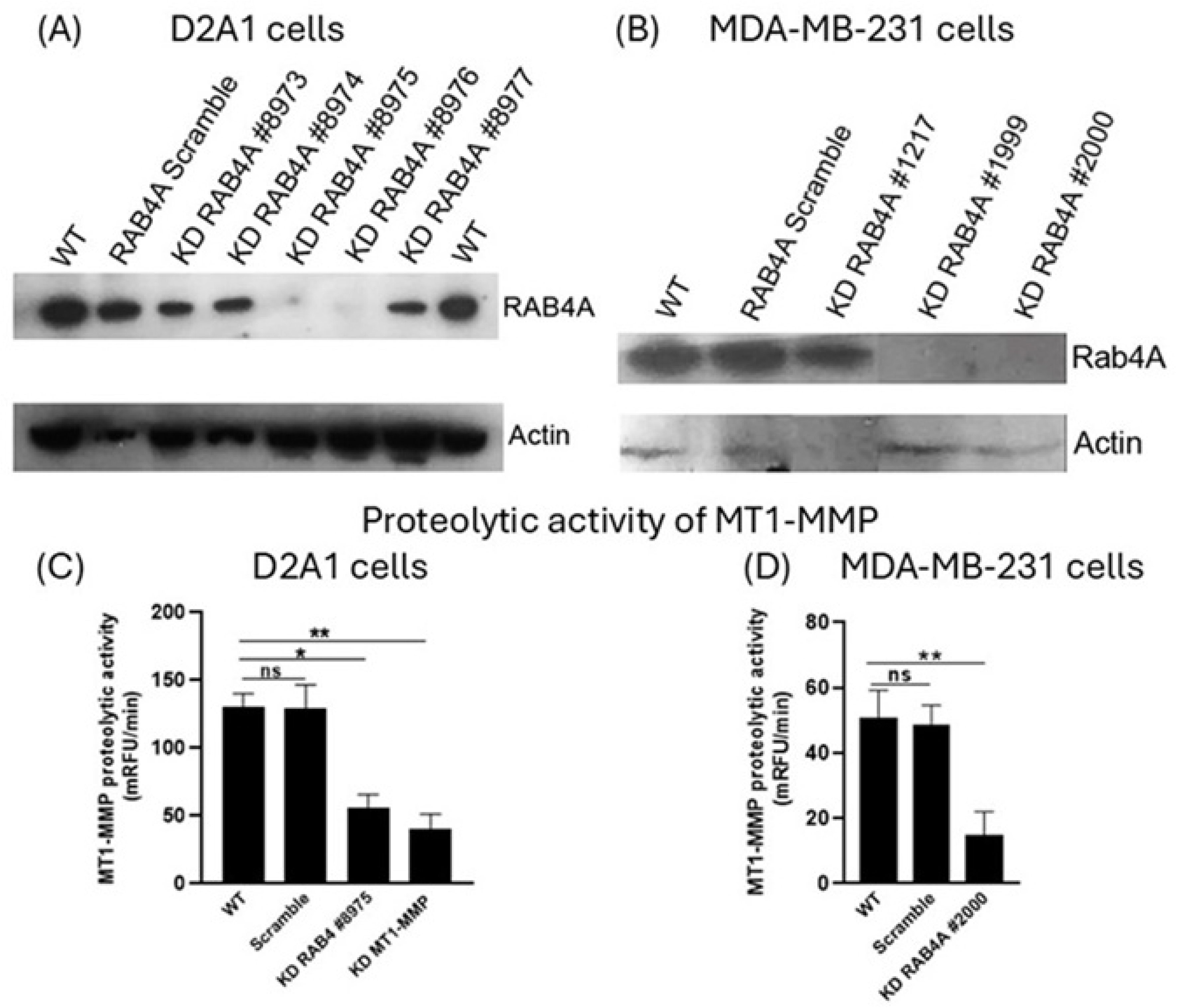
Downregulation of RAB4A reduces MT1-MMP proteolytic activity. Western blot illustrating the downregulation of RAB4A mediated by shRNA in (A) TNBC murine D2A1 cells and (B) human TNBC MDA-MB-231 cells. Downregulation of RAB4A decreased MT1-MMP proteolytic activity as determined with the fluorogenic peptide MMP-14 Substrate I. (C) A reduction by 2.3-fold was measured in the D2A1 KD RAB4A cells and (D) 3.4-fold in the MDA-MB-231 KD RAB4A cells. *p < 0.05, **p < 0.005.

A proteolytic assay with the fluorogenic probe MMP-14 Substrate I was performed to confirm that downregulating RABA4 reduces MT1-MMP activity at the cell surface (Figure 2C and D). As control, the proteolytic activity of MT1-MMP was measured in the D2A1 KD MT1-MMP cells. For these cells, we have previously reported a 70% reduction of MT1-MMP expression as measured by quantitative polymerase chain reaction (qPCR) and at the protein level by Western blot [21]. As expected, downregulation of the MT1-MMP gene decreased by 3.4-fold its proteolytic activity on the cell surface compared to D2A1-WT and D2A1 RAB4A-scramble cells (*p* < 0.005) (Figure 2 C). Downregulation of RAB4A led to a 2.3-fold reduction in MT1-MMP proteolytic activity in D2A1 KD RAB4A cell (*p* < 0.05), and of 3.4-fold reduction in MDA-MB-231 KD RAB4A cells (*p* < 0.005). The shRNA scramble sequence did not significantly alter RAB4A expression or MT1-MMP proteolytic activity in the D2A1 RAB4A-scramble cells and MDA-MB-231 RAB4A-scramble cells.

### Reduction of cancer cell invasion *in vitro* by downregulating RAB4A

Downregulation of RAB4A or MT1-MMP expression in D2A1 cells similarly reduced the invasive capacity by approximately 2.1-fold (*p* < 0.005), compared to the wild-type counterpart, when the plasma collected before radiotherapy was used as attractant (Figure 3A). A similar reduction was measured with the MDA-MB-231 KD RAB4A cells that have crossed 2.9-fold less the Matrigel layer of the invasion chambers than wild-type MDA-MB-231 cells (*p* < 0.0005), (Figure 3B). As a control, only a small but statistically significant reduction in invasion capacity was observed when these TNBC cell lines were transfected with the scrambled shRNA sequences (*p* < 0.005). In supplementary controls, 1% plasma collected before radiotherapy and 1% FBS used individually as attractant have increased similarly the invasiveness of MDA-MB-231 cells by 4.3-fold, compared to plasma-free (0.1% BSA) (Figure 3B).

**FIGURE 3.**
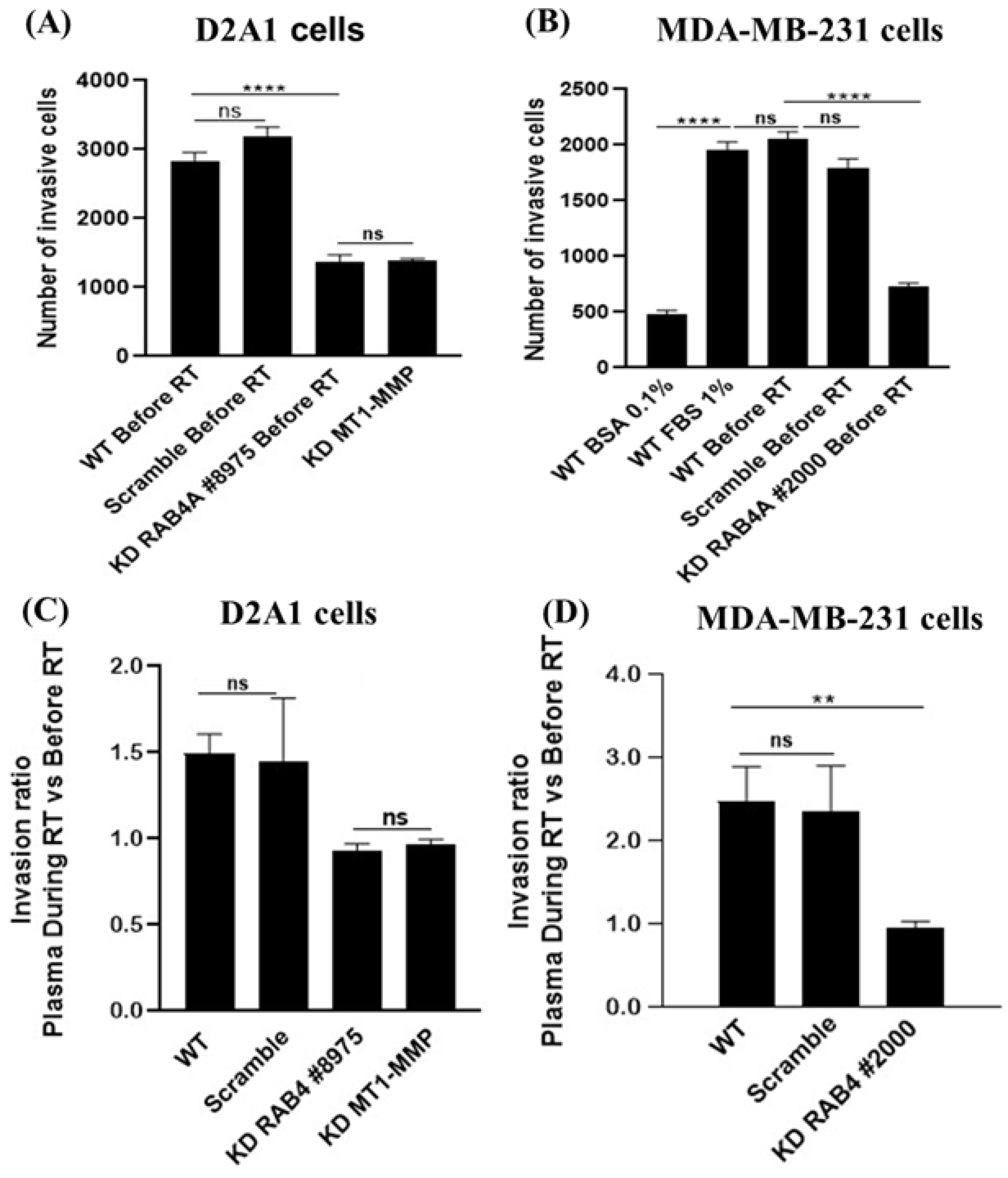
Invasion capacity of D2A1 and MDA-MB-231 cells. The downregulation of RAB4A and MT1-MMP resulted in a decreased invasion capacity of TNBC cells when plasma collected before radiotherapy (RT) was used as an attractant (A, B). Additionally, these downregulations significantly inhibited the stimulation of invasion caused by plasma collected during radiotherapy (C, D), highlighting the relevance of targeting these pathways. *p < 0.05, ***p < 0.005 and ****p < 0.0005.

### RAB4A downregulation prevents radiation-stimulation of cancer cell invasion

Plasmas collected from the TNBC patient before and during radiotherapy were added individually as attractant in the lower compartment of the invasion chambers. The results are reported according to the ratio of the number of cells having passed through the Matrigel layer with the plasma collected during the radiotherapy / plasma collected before the radiotherapy. For D2A1 WT cells and those transfected with the shRNA scramble sequence, plasma collected during radiotherapy increased their invasive capacity by approximately 1.6-fold (*p* < 0.0005) (Figure 3C). A larger stimulation was achieved with the MDA-MB-231 WT cells and its scramble clone whose invasion capacity was increased by 2.5-fold (*p* < 0.005) (Figure 3D). The radiation-induced stimulation of cancer cell invasion was largely blocked by downregulating RAB4A. Indeed, the same number of D2A1 KD RAB4A and MDA-MB-231 KD RAB4A cells crossed the Matrigel layer when incubated with plasma collected before or during radiotherapy. Supporting the role of MT1-MMP, its downregulation in D2A1 KD MT1-MMP cells also largely abolish the stimulatory effect of plasma collected during radiotherapy on cancer cell invasion (Figure 3D).

### Impact of RAB4A downregulation on the development of lung metastases

The D2A1-WT, D2A1 RAB4A-scramble and D2A1 KD RAB4A cells expressing the fluorescent protein FUCCI were incubated *in vitro* in DMEM medium supplemented with either 0.1% BSA, 1% plasma collected before radiotherapy, or 1% plasma collected during radiotherapy. Twenty-four hours later, they were washed and injected i.v. into tail of female Balb/c mice. The number of lung metastases were quantified 28 days later with the epifluorescence microscope EVOS FL Auto Imaging System.

Plasma collected before radiotherapy slightly increased the mean number of metastases, reaching 2.2 ± 1.4 per mouse, compared to 0.5 ± 0.5 for the control incubated with 0.1% BSA. A more significant amplification was observed with plasma collected during radiotherapy, increasing the mean number of metastases to 7.6 ± 2.9 (*p* < 0.05) (Figure 4).

**FIGURE 4.**
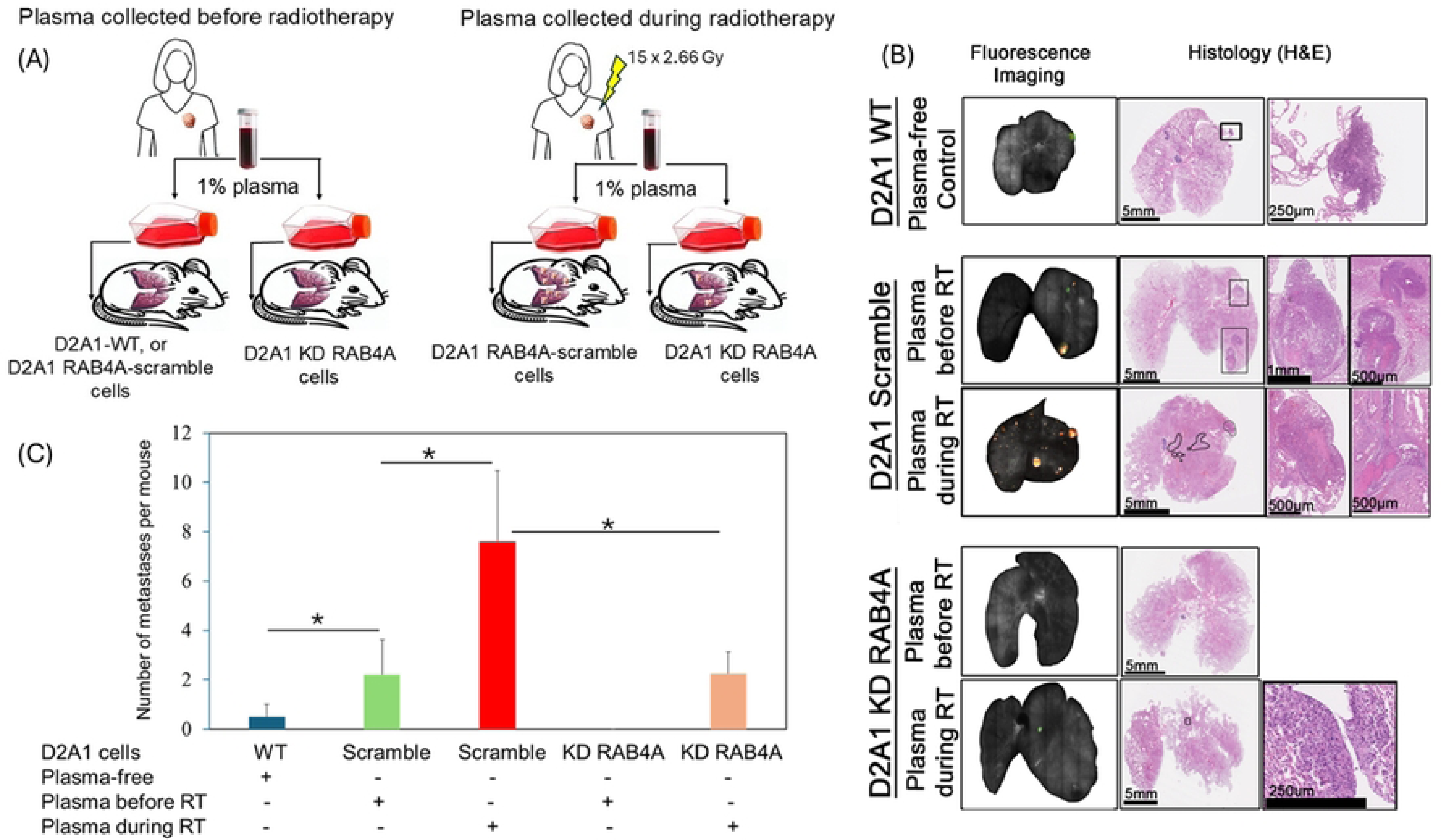
Downregulation of Rab4A prevents metastasis development induced by plasma collected from a TNBC patient during radiotherapy. (A) Plasma samples were collected before and during radiotherapy from a TNBC participant who relapsed within one year of treatment. These plasmas were then incubated with D2A1 TNBC cells and subsequently injected intravenously into Balb/c mice. (B) Lung metastases were quantified 28 days later using fluorescence imaging, and their presence wasconfirmed by histology. (C) Development of lung metastases by plasma collected during radiotherapy was averted by downregulating Rab4A. *p < 0.05.

RAB4A downregulation completely blocked metastasis development when D2A1 KD RAB4A cells were preincubated with plasma collected before radiotherapy. Interestingly, this protective effect was also obtained when plasma collected during radiotherapy was tested with the D2A1 RAB4A cells, resulting in a reduction of metastases to 2.3 ± 0.9, compared to 7.6 ± 2.9 when this plasma was incubated with the D2A1 RAB4A-scramble cells (*p* < 0.05).

## Discussion

The relapse rate in early-stage TNBC patients is significantly higher compared to the other breast cancer subtypes [3]. This observation highlights the need to identify new molecular targets to develop drugs capable of preventing the development of metastases which are detected after treatments. This study aims to evaluate the relevance of RAB4A as a potential target to achieve this objective.

Rab4A expression progressively increases from normal tissue to primary then to metastatic tumor [33]. Rab4A is amplified in various tumors, especially in invasive breast cancers [33, 34], and its high expression correlates with poorer prognosis [33, 34, 37]. Gathering evidence indicates that Rab4A is required for development of metastases [33, 34, 37–42]. Rab4A overexpression in MDA-MB-231 cells drives tumor cell invasive migration in 3D Matrigel assays [43]. Silencing of Rab4A or expression of a dominant-negative (Rab4DN) mutant in MDA-MB-231 and MCF10.DCIS.com breast cancer cells reduced MT1-MMP recycling, extracellular matrix degradation and invasion in 3D gel matrices, and importantly, inhibited cancer cell invasiveness in mammary gland and lung tissue in immunodeficient mice [34].

The current study suggests that Rab4A may also play a role in increasing the invasiveness of cancer cells and the development of metastasis triggered by radiotherapy in TNBC.

The development of metastases results from a multi-step process, including the invasion of cancer cells from the primary tumor into surrounding tissues, intravasation into the bloodstream, and extravasation to a distant organ where cancer cells colonize and grow [12]. A key molecular event is the cleavage of the extracellular matrix proteins by MMP, which creates a pathway for cancer cells to colonize distant organs. Their essential role in tumor progression has generated considerable interest in developing inhibitors to target them. However, convincing clinical results are still awaited because of the poor bioavailability and selectivity of MMP inhibitors [44–46]. One constraint to their clinical application could be the significant role of MMPs in fundamental physiological processes, such as promoting angiogenesis, cell migration, modulation of inflammation response and osteoclast activity [47].

MT1-MMP is also essential in numerous physiological processes [47]. However, its localization on the surface of cancer cells could offer an advantage in managing cancer progression. Over the past decade, inhibitors of MT1-MMP have been developed [48]. Although they can bind to the active sites of MT1-MMP, their overall effectiveness has remained limited [44]. The past failures in clinical trials have been mainly associated to adverse effects. Waiting for more MT1-MMP-specific inhibitors [48], we propose a novel approach which is to reduce MT1-MMP trafficking on the surface of cancer cells by inhibiting RAB4A.

To assess the potential of this new opportunity, expression of RAB4A have been downregulated by shRNA in the D2A1 and MDA-MB-231 cells. The reduction of MT1-MMP on surface of these cancer cell surface was confirmed by a significantly decreased of its proteolytic activity, resulting in a marked loss of invasion capacity *in vitro* for the D2A1 and MDA-MB-231 cells. Downregulation of RAB4A in D2A1 tumor implanted in a Balb/c mouse mammary gland also completely prevented the spontaneous development of lung metastases.

Although this approach is promising, RAB4A inhibition cannot be performed in disease-free individuals to prevent the development of metastases because MT1-MMP is involved in multiple physiological processes. A RAB4A inhibitor should therefore be administered to eliminate metastases or to prevent their development in early-stage TNBC patients who are initially diagnosed as free of metastases.

Emerging evidence indicates that the inflammatory response triggered by radiotherapy may promote critical pathways in certain patients, such as increased cancer cell invasion, a higher number of circulating tumor cells, and the development of new metastases [20, 49–51]. Ongoing clinical trials will provide further results to help understand the association between radiotherapy and the increased risk of developing metastases for certain patient groups. In the meantime, it is relevant to evaluate whether an MT1-MMP inhibitor could block radiotherapy stimulation of metastasis development.

In this study, a TNBC patient was recruited to determine whether downregulating RAB4A might block radiotherapy stimulation of some critical steps in cancer progression. Plasma samples were collected before and after the 4^th^ fraction of radiotherapy in a patient with an early-stage TNBC (cT2N0M0). At diagnosis, the absence of metastases was confirmed by positron emission tomography (PET). Six months after radiotherapy, metastases in lung, bone and brain were detected.

The same number of MDA-MB-231 cells has migrated through the Matrigel layer of the invasion chamber when 1% FBS or 1% plasma collected before radiotherapy were used as attractant. On the other hand, 1% plasma collected during radiotherapy significantly enhanced the invasiveness of the TNBC cell lines MDA-MB-231 and D2A1. Additionally, a pre-incubation *in vitro* of the D2A1 cells followed by their injection i.v. into tail vein of mice markedly promote the development of lung metastases. This TNBC patient appears to belong to a subgroup where early recurrence is linked to an increased ability of cancer cells to invade and metastasize after radiotherapy Therefore, she was a suitable candidate to assess whether downregulating RAB4A could help to block these side effects of radiotherapy. Our preclinical data clearly show the downregulating RAB4A markedly reduces radiotherapy stimulation of cancer cell invasion and metastasis development. These results were expected since similar protective effects were reported in preclinical models when MT1-MMP was downregulated [21, 28].

## Conclusions

Reducing the relapse rate occurring within the first 3 years following treatment for early-stage TNBC patients still represents a significant challenge in improving the management of these patients. Since radiotherapy remains an essential component in the treatment options for TNBC, the next step is to develop a complementary treatment that will reduce the incidence of recurrence that could be associated with radiotherapy. Among the targets that deserve to be exploited, inhibition of the ability of RAB4A to recycle MT1-MMP on the surface of cancer cells could prove promising. Along with developing this inhibitor, it will be necessary to determine the period of its administration before, during and after the radiotherapy plan to ensure that patients receive benefits while maintaining good tolerance.

## ACKNOWLEDGMENTS

This work was supported by Canadian Institutes of Health Research, grant number 418761, obtained by the authors J.L.P. and B.P. The funders had no role in study design, data collection and analysis, decision to publish, or preparation of the manuscript

